# Deciphering complex metabolite mixtures by unsupervised and supervised substructure discovery and semi-automated annotation from MS/MS spectra

**DOI:** 10.1101/491506

**Authors:** Simon Rogers, Cher Wei Ong, Joe Wandy, Madeleine Ernst, Lars Ridder, Justin J.J. van der Hooft

## Abstract

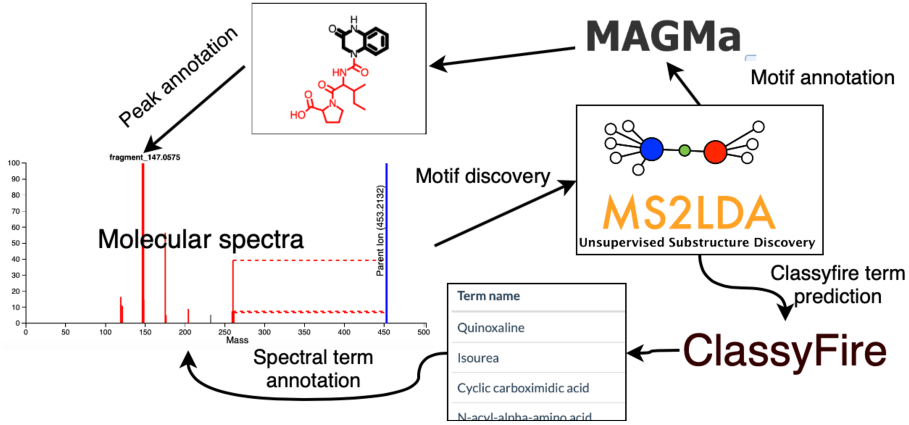
Integration of MS2LDA substructure discovery with MAGMa spectral annotations and ClassyFire term predictions complemented with MotifDB significantly advance metabolite annotation.

Complex metabolite mixtures are challenging to unravel. Mass spectrometry (MS) is a widely used and sensitive technique to obtain structural information on complex mixtures. However, just knowing the molecular masses of the mixture’s constituents is almost always insufficient for confident assignment of the associated chemical structures. Structural information can be augmented through MS fragmentation experiments whereby detected metabolites are fragmented giving rise to MS/MS spectra. However, how can we maximize the structural information we gain from fragmentation spectra?

We recently proposed a substructure-based strategy to enhance metabolite annotation for complex mixtures by considering metabolites as the sum of (bio)chemically relevant moieties that we can detect through mass spectrometry fragmentation approaches. Our MS2LDA tool allows us to discover - unsupervised - groups of mass fragments and/or neutral losses termed Mass2Motifs that often correspond to substructures. After manual annotation, these Mass2Motifs can be used in subsequent MS2LDA analyses of new datasets, thereby providing structural annotations for many molecules that are not present in spectral databases.

Here, we describe how additional strategies, taking advantage of i) combinatorial *in-silico* matching of experimental mass features to substructures of candidate molecules, and ii) automated machine learning classification of molecules, can facilitate semi-automated annotation of substructures. We show how our approach accelerates the Mass2Motif annotation process and therefore broadens the chemical space spanned by characterized motifs. Our machine learning model used to classify fragmentation spectra learns the relationships between fragment spectra and chemical features. Classification prediction on these features can be aggregated for all molecules that contribute to a particular Mass2Motif and guide Mass2Motif annotations.

To make annotated Mass2Motifs available to the community, we also present motifDB: an open database of Mass2Motifs that can be browsed and accessed programmatically through an API. MotifDB is integrated within ms2lda.org, allowing users to efficiently search for characterized motifs in their own experiments. We expect that with an increasing number of Mass2Motif annotations available through a growing database we can more quickly gain insight in the constituents of complex mixtures. That will allow prioritization towards novel or unexpected chemistries and faster recognition of known biochemical building blocks.

## Introduction

Complex natural mixtures are full of specialized metabolites with diverse structures and functions.^1^ In untargeted metabolomics approaches these molecules give rise to information-rich mass spectral data sets and a key challenge is the interpretation of this data, particularly in terms of identifying chemical structures.^2, 3^ This process is commonly referred to as metabolite annotation and identification,^4^ a highly challenging process which typically enables the assignment of chemical structures to only a very small percentage of molecules detected.^2, 5-7^ Consequently, the rapid and automated identification of chemical structures from complex mixtures is one of the main obstacles hindering for example the discovery of novel bioactive molecules addressing global health care threats such as antimicrobial resistance, cancer or inflammatory diseases.

Recently, we have introduced the concept of Mass2Motifs, and the MS2LDA approach to find these molecular substructure-related mass fragmentation patterns in mass spectral metabolomics data.^8^ We showed that through Mass2Motif discovery, we can assign substructures to more than 70% of the fragmented molecules in beer extracts. Another widely used tool to organize fragmentation spectra is mass spectral Molecular Networking.^9, 10^ In combination or as stand-alone tool, these MS2 spectral similarity-based grouping algorithms are the current state-of-the-art in the untargeted metabolomics field to rapidly get a comprehensive overview of molecular diversity in samples.^11-15^ To retrieve chemical structural information for acquired experimental spectra, MS2 fragmentation patterns are matched directly to library reference data or *in silico* by matching substructures of candidate structures,^5, 16, 17^ however only a very low percentage of the molecular features (typically 2-5% up to 30% in rare cases) can be confidently assigned to known chemical structures. In comparison to the structural annotation of entire molecules, structural annotation of the Mass2Motifs is more straightforward and less complex, as Mass2Motifs represent smaller substructures. However, the structural annotation of Mass2Motifs is currently performed by a combination of manual peak search in MS/MS databases such as MetLin^18^ and MzCloud^19^ as well as expert knowledge, and thus still represents a tedious and time-consuming step, especially for large-scale high-throughput experiments with several hundred discovered Mass2Motifs per experiment. As we and others have shown, ^8, 17, 20, 21^ the use of reference MS/MS spectra of standards speeds up the annotation process; however, with the increasing size of publicly available MS/MS reference libraries, ^9, 17^ complete manual Mass2Motif annotation and curation is rapidly becoming impractical. Furthermore, with the expected increase in publicly available experimental MS/MS data the amount of structurally novel Mass2Motifs is expected to steadily rise. This will make structural predictions for Mass2Motifs of non-standards and effective reuse of previously annotated Mass2Motifs essential. Thus, the next step is to semi-automate Mass2Motif annotation and store annotated Mass2Motifs such that they can be used in the future. Here, we propose and implement two complementary strategies for semi-automated motif annotation and introduce MotifDB for Mass2Motif reuse.

In recent years, algorithms that propose chemical substructures and candidate structures for mass features have become available.^22-25^ For example, MAGMa maps possible candidate molecules to MS/MS spectra in experimental data by assigning possible substructures from a candidate molecule to the mass fragments and subsequently ranks different candidate molecules using those annotations based on a relatively simple scoring algorithm.^26^ Here, we used MAGMa for automated annotation of the features within Mass2Motifs based on the MAGMa annotations of reference spectra, using the known chemical structures as candidates. The results of this MAGMa-MS2LDA integration are made available via the ms2lda.org web app, and can help the annotation of new motifs and may support users to chemically interpret the presence of motifs in unknown molecules.

A complementary strategy towards structural annotation is to predict molecular properties such as fingerprints or classification based on spectral features.^27, 28^ To annotate Mass2Motifs, one would need to learn and combine such molecular properties for the Mass2Motif MS/MS features. If MS2LDA is applied to a set of reference molecules, chemical class predictions of Mass2Motifs can be performed by augmenting molecular properties directly from their known structures. Here, we stored ClassyFire^29^ substituent terms for each standard and determined enriched class terms upon comparing the presence of those class terms within a Mass2Motif versus the entire data set.

Prediction of chemical properties for Mass2Motifs discovered in non-standards from the associated MS/MS spectra is a more challenging task. Here, we decouple this into two separate tasks (collectively known as ClassyFirePredict), i) training a model that links ClassyFire substituent terms of molecules to MS/MS spectral features using reference data, and ii) analyzing enriched substituents/fingerprints within Mass2Motifs from non-standards data by applying the trained model from i). Using a publicly available annotated MS2LDA experiment, we show how this can guide the user for annotation of fragment-based Mass2Motifs such as flavonoid and saccharide related motifs.

Finally, to effectively reuse previously annotated motifs, we introduce MotifDB that is available from the ms2lda.org web app.^30^ Currently, the MotifDB serves as a result for annotated Mass2Motifs with their MS/MS features. A number of annotated Mass2Motif sets from various source including plant extracts, urine, and standards, are already available for matching against Mass2Motifs discovered in new experiments. The extensions to the original ms2lda.org platform presented here are shown schematically in Figure 1.

**Figure 1:**
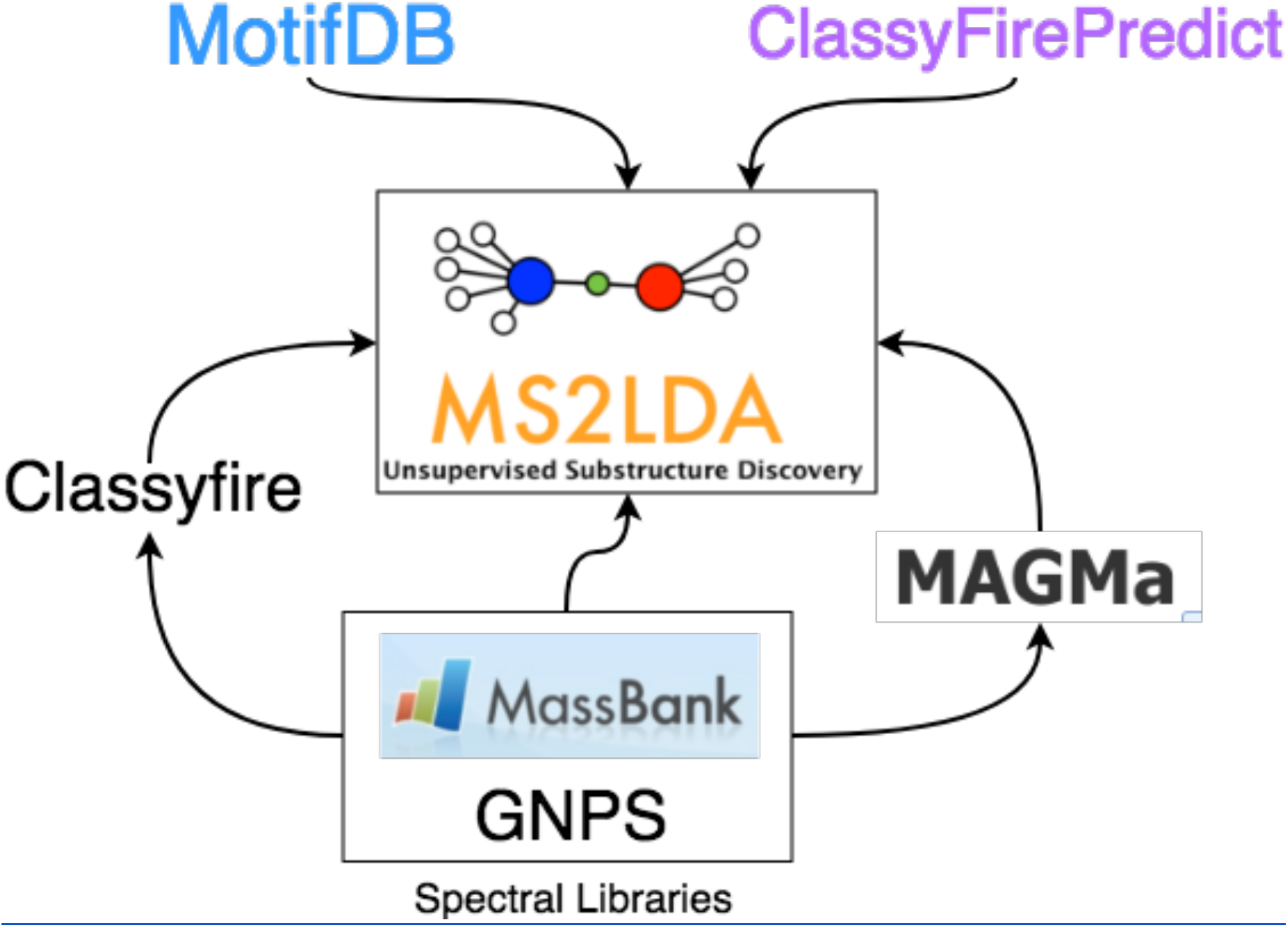
The extensions to the original MS2LDA model described in this paper. MotifDB provides a platform for storing annotated Mass2Motifs. MAGMa and Classyfire are both used with standard datasets to provide insight into the structural makeup of the MassMotifs derived from them. ClassyFirePredict extends this idea to non-standard data by predicting Classyfire terms directly from the mass spectra.

We expect that the here described augmentations to the ms2lda.org web app will empower researchers with the tools to more rapidly decipher complex mixtures and create annotated and curated sets of Mass2Motifs. Those in turn will be effective in future experiments to more quickly assess the presence of specific molecular types in complex mixtures and assess chemical diversity of those mixtures based on substructure recognition. We expect these substructure-based annotation strategies to become essential to decipher complex mixtures and enable meaningful biochemical interpretation.

## Methods

### Fragment spectrum to ClassyFire substituent term classifier for standards and non-standards + neural network for classification of Mass2Motifs from non-standards

ClassyFire terms were derived through the CLassyFire API for two of the public standard datasets (massbank_binned_005 and gnps_binned_005 - see Data Availability section) stored within ms2lda.org using the ClassyFire web API^29^ based on the molecules’ InCHiKeys. The substituent terms were stored in the database and linked to the relevant molecules such that they are visible when the molecule is explored. Additional functionality was added to ms2lda.org to summarize the terms within a particular Mass2Motif. In particular, based on actual values of the fragment spectra to Mass2Motif probability and overlap score thresholds as outputted by MS2LDA,^31^ the molecules associated with each Mass2Motif are extracted, along with their ClassyFire substituent terms. For each term, the proportion of molecules associated with the Mass2Motif that include the term is computed, along with the proportion of molecules in the experiment. Comparing these terms provides evidence as to how unique and concentrated that term is in the Mass2Motif.

When working with new experimental data, exploring ClassyFire terms from standard molecules is useful if a discovered motif closely matches one of those in the standards experiments. To further extend this functionality we have developed a machine learning approach that can predict putative ClassyFire terms from any mass spectrum. A multilayer neural network was produced that, for a binned mass spectrum, predicts the probability of presence/absence for each ClassyFire term. The network was built in Python using keras.^32^ Spectral data are currently binned into bins of width 1Da with *m/z* values over 1000 discarded. After normalizing so that the base bin (i.e. the bin holding the most intensity in a particular spectrum) had intensity of 1000.0, the data were log transformed (after adding 1.0 to avoid problems associated with taking the log of zero). The network consists of a 1000-dimensional densely-connected input layer, followed by two hidden dense layers (of dimension 500 and 200) and then an output layer with dimension equal to the number of ClassyFire substituent terms. Non-linear ReLU (rectified linear unit) activation functions were used for the hidden layers, and a sigmoid function for the output layer. The model was optimized using the binary cross entropy loss function. This model represents our initial network design and it is likely that it could be optimized further.

An initial training and validation phase was used to determine which terms could be reliably predicted. A filtered dataset of 10,038 unique tandem mass spectra with associated chemical structures retrieved from the Global Natural Products Social Molecular Networking (GNPS) platform was used that was created as follows. Mass spectral molecular networks were created using all publicly available libraries on GNPS and these were matched back to the libraries to obtain chemical structural information. The resulting mass spectral molecular networks and results from spectral library matching are publicly available (see Data Availability section). Subsequently, we used a script in python (see Code availability section) to sub-select only tandem mass spectra with associated chemical structural information on their complete structure in computer readable format to create a dataset in the .MGF data format followed by the selection of 10,105 unique molecules based on the first 14 digits of the InchiKeys with parent *m/z* < 1000 Da. From these selected molecules, we could get classifications from the ClassyFire API for 10,038 of them resulting in the final data set.

Ten random splits into training (90%) and validation (10%) were used to assess the performance with respect to each term. Within each split, the area under the receiver operating characteristic curve (AUC) was computed, and these were averaged across the ten splits. Figure 2 describes the distribution of AUCs for the 2098 unique terms found in the dataset. Note that AUCs of less than 0.5 (worse than random guessing) occur due to the very small number of occurrences of some terms (1804 of the 2098 terms appear in fewer than 1% (100) of the 10,038 unique molecules). Based on this analysis, we selected 444 terms for the final classifier as including terms that were proven to be unpredictable would only add noise to the ms2lda.org analysis. These 444 terms were chosen via two conditions. Firstly, all terms with an average AUC across the ten splits of greater than 0.7 and also, terms with an AUC of between 0.6 and 0.7 who appeared in at least 0.5% of the molecules in the dataset. These additional terms were included to increase coverage under the assumption that some false positives can be tolerated for individual molecules as they are likely to be filtered out when we explore terms at the Mass2Motif level. Finally, the model was re-trained using these 444 terms and all of the available training data.

**Figure 2:**
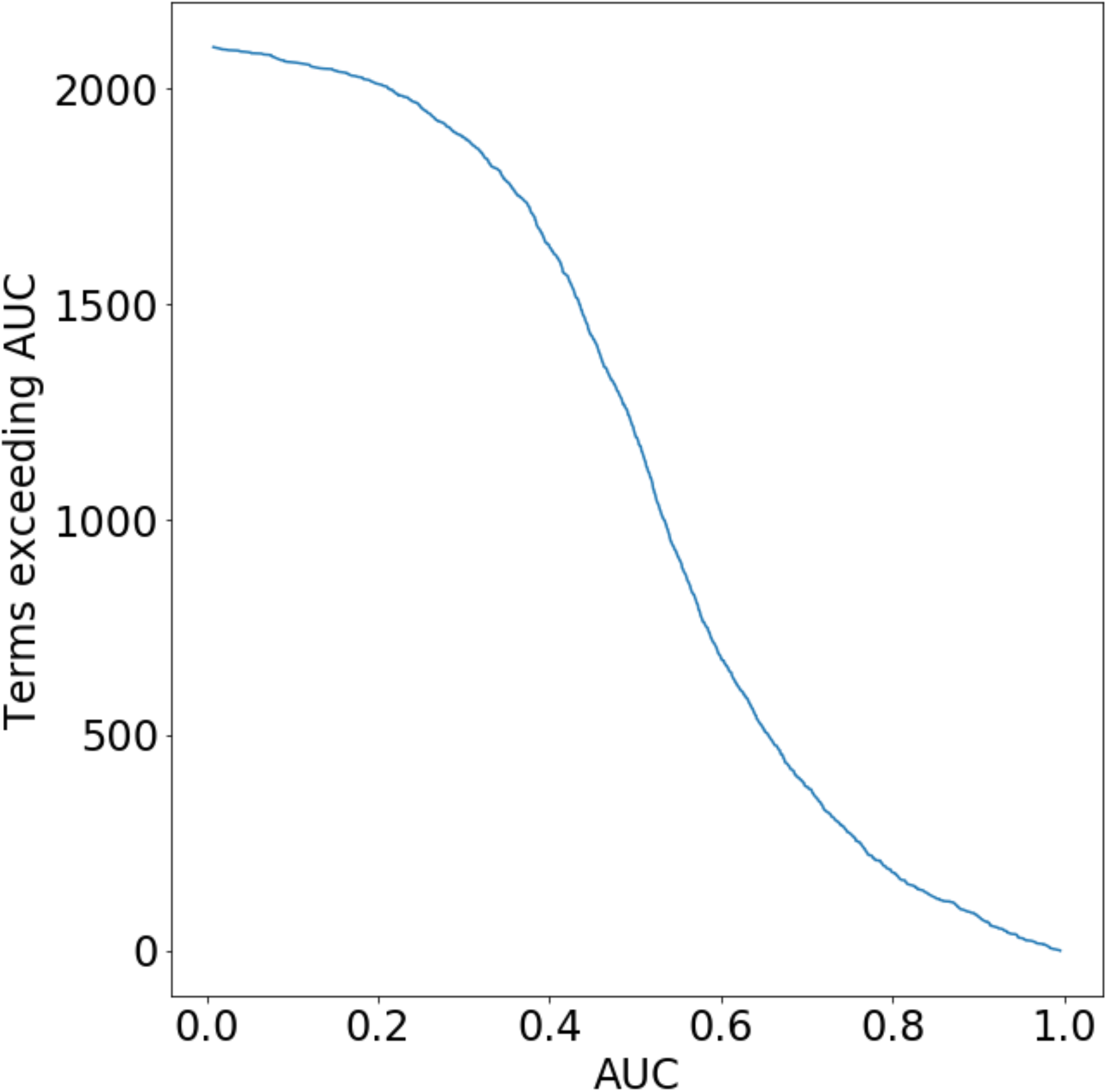
The number of terms exceeding different AUC values (y-axis) versus the AUC (x-axis). The curve shows the number of terms that exceed any particular AUC value. Note that AUCs of less than 0.5 (worse than random guessing) occur due to the very small number of occurrences of some terms (1804 of the 2098 terms appear in fewer than 1% (100) of the 10,038 unique molecules). Terms with AUCs above 0.7 and those between 0.6 and 0.7 appearing in at least 0.5% of the molecules in the dataset were used to train a neural network.

The predictive model was incorporated into ms2lda.org allowing users to assign putative ClassyFire terms to any molecules. These terms are then collated at the Mass2Motif level to aid in annotation in exactly the same manner as those linked to the reference molecules.

### MAGMa-MS2LDA integration

We used MAGMa to annotate Mass2Motif features as follows. All reference spectra for four data sets of known molecules that were subjected to MS2LDA (massbank_binned_005, gnps_binned_005, 2613 public spectra from various sources in positive ionization mode, and 551 public spectra in negative ionization mode from various sources - see Data Availability section), were analyzed and annotated using MAGMa (see Code Availability section). The annotation of each spectrum was performed with a single candidate molecule, i.e. the known molecule. As a result, the likely substructures of this molecule were assigned to individual peaks in the spectrum. Note, that only the peaks used in the MS2LDA analysis were included in the MAGMa analysis and that not all of those necessarily match with a simple substructure found within the reference molecule. Subsequently, the substructures were matched with the actual features used in the MS2LDA analysis (either fragments or losses within user-defined mass bins). For fragment features, the substructures assigned by MAGMa were stored both as a canonical SMILES, generated by RDKit, and as a mapping (with atom indices) on the original molecule. A SMILES string was generated for the loss features by first removing the MAGMa substructure atoms from the parent molecule and generating a canonical SMILES from the remaining atoms. These SMILES may contain disconnected parts of the molecules (separated by a dot according the SMILES specifications). These MAGMa substructure feature annotations were stored in the database of MS2LDA, and visualized on the web interface where substructures are drawn using the ChemDoodle package.^33^

As a result, Mass2Motif pages in MS2LDA could now be augmented with the MAGMa substructure annotations as follows. For a given feature explained by a Mass2Motif, all substructures associated to the feature in the corresponding spectra (MS2LDA documents) are retrieved and grouped based on the canonical SMILES. Note that the same feature in different spectra may correspond to different substructures. In the current interface, all unique substructures are presented together with the number of times they occur in the corresponding spectra. Additionally, since the same binned fragment and neutral losses are used as global features across all experiments in MS2LDA.org, annotations for all (and new) features that have corresponding features in MAGMa-annotated experiments can be derived from the existing MAGMa annotations assigned to these shared global features. We show this new information in the Mass2Motif and Document pages of the ms2lda.org web app.

### MotifDB

Once Mass2Motifs have been annotated, it is useful to be able to search for them in future MS2LDA experiments. To this end, we have created a new application within ms2lda.org called MotifDB: a database for annotated Mass2Motifs (http://ms2lda.org/motifdb). This database can be accessed via an Application Programming Interface (API) as well as being searchable against other experiments in the ms2lda.org web app. In particular, when an experiment has been run through MS2LDA.org, the user can start a motif matching procedure against MotifDB whereby the Mass2Motifs discovered in the user’s experiment are compared against those in the MotifDB. Where a match exceeds a user-defined threshold (matches are computed via a cosine similarity score), the experimental motif can be linked to the MotifDB motif, which in turn will transfer the MotifDB annotation and allow it to be highlighted in visualizations. As MotifDB grows by community efforts, more and more Mass2Motifs learnt in experiments will be available to be matched with a high score against those in the MotifDB database, allowing for more rapid characterization of diverse chemical mixtures.

### Code Availability

The python script to generate MAGMa annotations of standards datasets is provided on github: https://github.com/iomega/motif_annotation

The python script to collect all GNPS library compounds is provided on github: https://github.com/madeleineernst/EditMGF/blob/master/CompileGNPSMGF_withInChIKey.py

The scripts to prepare the GNPS library molecules for neural networking and perform the neural networking is provided on github: https://github.com/sdrogers/nnpredict

Code to perform MS2LDA is available at: https://github.com/sdrogers/lda

Code for the ms2lda.org visualisation platform is available at: https://github.com/sdrogers/ms2ldaviz

### Data Availability

The following GNPS experiments were used in this manuscript:

Mass spectral molecular networks of public GNPS Library molecules - https://gnps.ucsd.edu/ProteoSAFe/status.jsp?task=6e22f85aeb0744208e872d1640f508d9

Library matching of the mass spectral molecular networks of GNPS Library molecules to retrieve full metadata - https://gnps.ucsd.edu/ProteoSAFe/status.jsp?task=03fba62d93cb4cbfa3f72106d18f7d2c

The following public MS2LDA experiments were used in this manuscript:

Reference molecules data sets:

massbank_binned_005 - http://ms2lda.org/basicviz/show_docs/190/

Gnps_binned_005 - http://ms2lda.org/basicviz/show_docs/191/

2613 public spectra from various sources in positive ionization mode - http://ms2lda.org/basicviz/summary/304/

551 public spectra in negative ionization mode from various sources - http://ms2lda.org/basicviz/summary/305/

Complex mixtures:

Urine38_POS_mzML_standardLDA_005binned - http://ms2lda.org/basicviz/summary/709

UrineDrugs_MolNetw_WorkshopSeattle2018 - http://ms2lda.org/basicviz/summary/601/

Rhamnaceae_plant_extracts_KyoBin_200Motifs_MS1_peaktable - http://ms2lda.org/basicviz/summary/566/

## Results

### MAGMa-based annotation of Mass2Motifs

#### MAGMa-MS2LDA annotations for previously analyzed Mass2Motifs

The integration of MAGMa with MS2LDA resulted in reference MS/MS MS2LDA experiments enriched with available MAGMa annotations for mass fragments and neutral losses for each fragmented molecule (Figure 3A). To evaluate how well these annotations matched with previously manually annotated motifs,^8^ we compared MAGMa annotations to those of manual annotations. These manual annotations were previously manually validated to be present or absent in motif-associated molecules which resulted in AUCs of around 0.69 for the GNPS and MassBank motif sets - thus indicating that within motifs the majority of the molecules contained the annotated substructures but a number of molecules contain unrelated substructures to those manual annotations. For example, motif 59 in the GNPS reference set was found to be related to the [Phenylalanine minus CHOOH fragment] substructure (http://ms2lda.org/basicviz/view_parents/58316/). Indeed, for 79 out of 117 molecules exactly this substructure was annotated by MAGMa for mass fragment 120.0825, with confirmation for the related aromatic fragment 103.0525 for 29 out of 35 appearances. This indicates that indeed this motif is related to [Phenylalanine minus CHOOH]; moreover, the MAGMa annotations also provide quick insight in structurally less related molecules in the motif that are included due to isomeric fragments giving rise to the same mass fragment. This highlights the need for manual validation of fragmentation patterns in molecules, which is now supported in the ms2lda.org web application.

**Figure 3:**
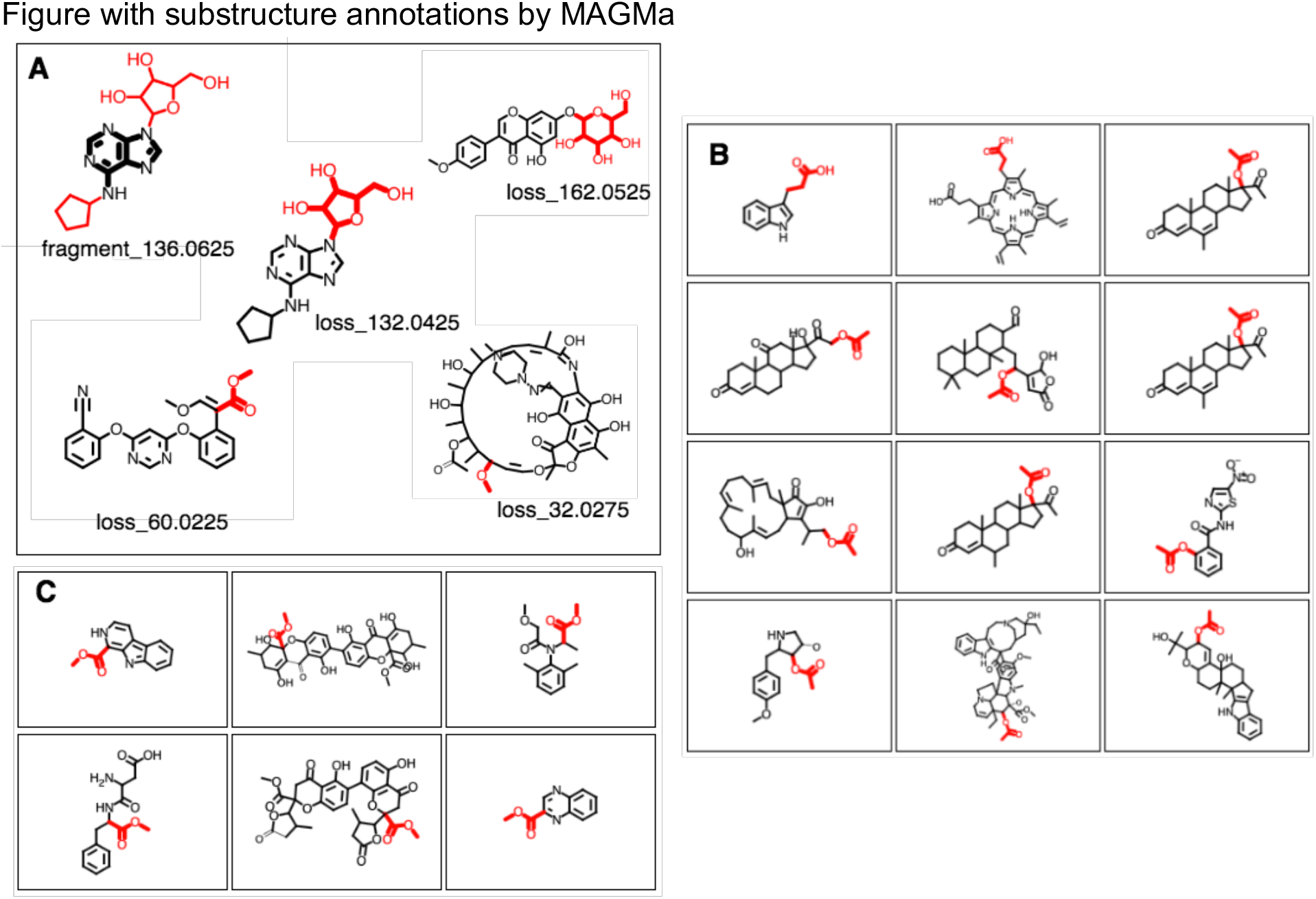
A-C Screenshots of the ms2lda.org web app with A) MAGMa annotations of Mass2Motif features in 5 motifs discussed in the results section. Annotated fragments are highlighted in black and bold, whereas annotated losses are depicted in red and bold. B) 12 examples of the 38 molecules for which the loss_60.0225 in GNPS Mass2Motif 49 was annotated with loss (CC(=O)O in SMILES. C) 6 examples of the 25 molecules for which the structurally related COC=O loss in SMILES was annotated for the same loss feature in GNPS Mass2Motif 49.

Another example is the indole related GNPS motif 25 (http://ms2lda.org/basicviz/view_parents/58017/); here, for 47 out of 110 molecules MAGMa annotated the 130.0675 mass fragment with a methylindole substructure and for 11 out of 28 molecules, the 118.0675 mass fragment was annotated with the indole substructure. Interestingly, the MAGMa annotations facilitated insight in other isomeric substructures within this motif; for example, MAGMa annotated the 130.0675 fragment for 17 molecules with a 2-aminopropyl-phenyl substructure and for 6 molecules the related 2-aminoethyl-phenyl substructure, indicating that motif 25 is also associated to this aromatic substructure. Other annotations for the 130.0675 fragment included two isobaric substructures with a different elemental formula which mass fell within the 0.005 Da mass bin.

MAGMa also annotated neutral loss-based Mass2Motifs. For example, GNPS Mass2Motif 49 which was annotated with “Loss possibly indicative of carboxylic acid group with 1-carbon attached” http://ms2lda.org/basicviz/view_parents/58174/. MAGMa annotations indeed show that the annotated loss (CC(=O)O in SMILES) was annotated to the largest set of 38 molecules out of 132 molecules as can be seen for 12 different molecules in Figure 3B. However, additional annotations were present such as 25 molecules containing the structurally related COC=O loss (Figure 3C). The remaining molecules contained isomeric losses. Similarly, an alternative loss was observed by MAGMa for GNPS motif 18 http://ms2lda.org/basicviz/view_parents/58383/ annotated as acetyl loss, as can be seen here: http://ms2lda.org/basicviz/show_doc/273058/. Furthermore, for Massbank Mass2Motif 41, “Loss indicative of [hexose minus H20]” it became clear that by far the majority of the MAGMa-annotated losses (50 out of 64) were glucose related http://ms2lda.org/basicviz/view_parents/57676/ (Figure 4A) with 13 being deoxyhexose moieties (Figure 4B) that - unlike usually - included the connecting oxygen atom upon fragmentation of the main scaffold. Similarly, with GNPS Mass2Motif 44, “[Pentose (C5-sugar)-H2O] related loss – indicative for conjugated pentose sugar”, for about half of the molecules (27 out of 56 molecules) the pentose loss was confirmed by MAGMa (Figure 4C) http://ms2lda.org/basicviz/view_parents/58179/. For this motif, alternative loss annotations were also annotated by MAGMa such as displayed in Figure 4D.

**Figure 4:**
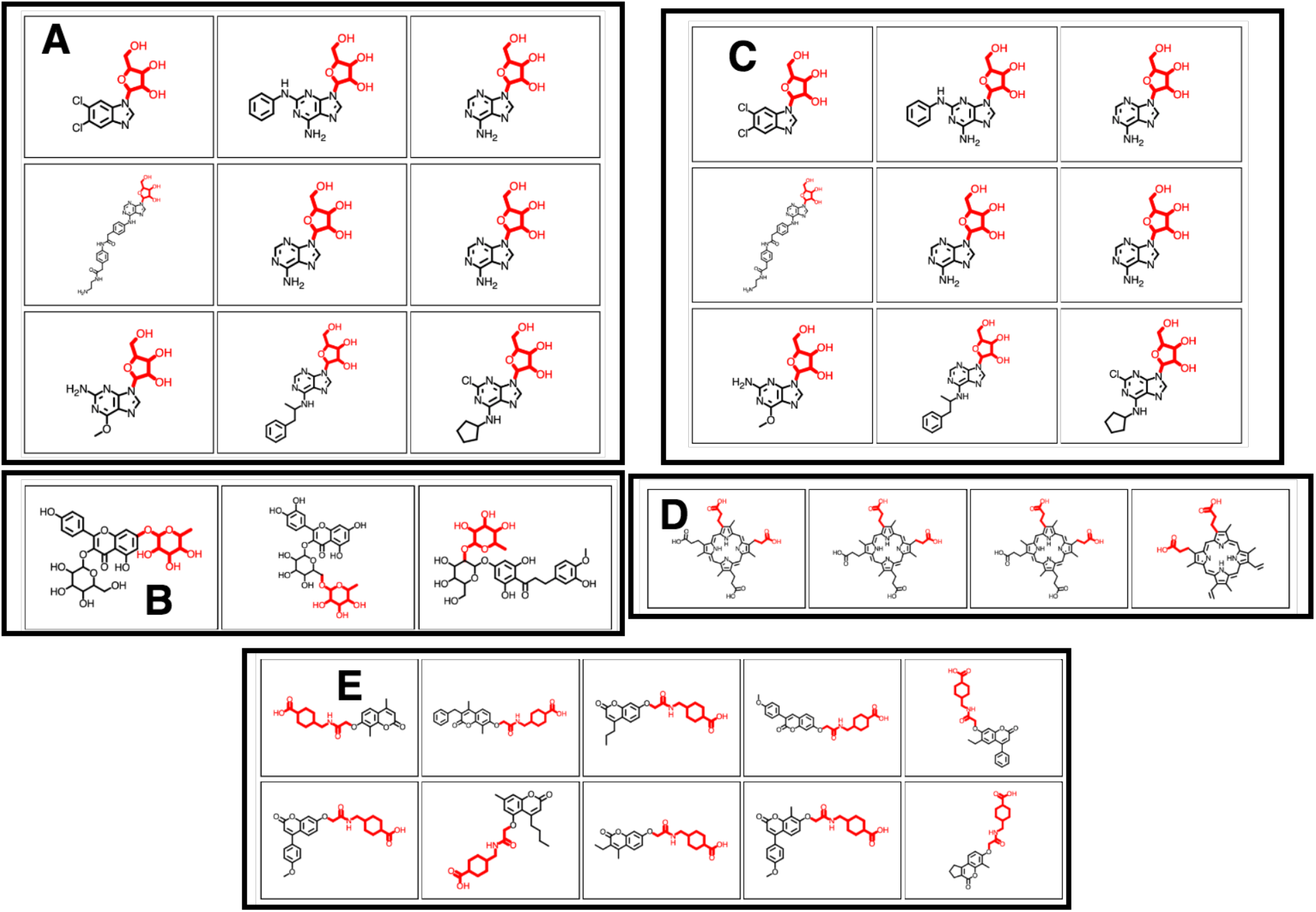
A-E Screenshots of the ms2lda.org web app with A) 9 different molecules out of the 50 molecules that MAGMa annotated with a hexose moiety for the loss feature in MassBank Mass2Motif 41. B) 3 examples of the 13 molecules where MAGMa annotated the loss feature in MassBank Mass2Motif 41 with a deoxyhexose moiety. C) 9 out of the 27 molecules for which MAGMa annotated a pentose moiety for the loss feature in GNPS Mass2Motif 44. D) alternative loss annotation of the loss feature in GNPS Mass2Motif 44. E) oxyacetyl-amino-methyl-cyclohexane-1-carboxylic acid loss annotated in 10 molecules of GNPS Mass2Motif 439.

Finally, GNPS motif 54 was annotated as ferulic acid related http://ms2lda.org/basicviz/view_parents/58325/. The MAGMa annotations show how important it is for this motif that the four mass fragments are all present since 73 molecules contained mass fragment 177.0525 whereas for mass fragment 117.0325 14 out of 19 molecules contained ferulic acid related substructures. Thus, whereas all GNPS Mass2Motif 54 related fragments have isomeric substructures unrelated to ferulic acid, their combined presence is highly indicative for the presence of ferulic acid.

#### MAGMa-MS2LDA integration for annotation of yet unexplored Mass2Motifs

In addition to previously annotated motifs, MAGMa annotations of not yet explored Mass2Motifs were analyzed. Figure 3A shows MAGMa annotations for Mass2Motif fragment and loss features for five of the here described motifs in one of their related molecules. For example, GNPS Mass2Motif 152 could now be easily annotated as methanol loss resulting from the presence of a methoxy group http://ms2lda.org/basicviz/view_parents/58033/. The methoxy related loss could be annotated in 51 out of 58 molecules by MAGMa. Another methoxy group related GNPS Mass2Motif (374) was uncovered where the loss of 16.0325 was assigned with CH4 in 33 out the 38 molecules in the motif. In addition, GNPS Mass2Motif 188 could be annotated as related to a 2-dimethylamine-ethanol loss *(m/z* 89.0825) which was present in 9 out of the 14 molecules http://ms2lda.org/basicviz/view_parents/58098/. Other examples where MAGMa facilitated motif annotations include MassBank Mass2Motif 315 (benzyl and phenoxy group containing molecules) where for 77 out of the 84 associated molecules the benzyl moiety was annotated by MAGMa. Moreover, in 20 molecules the phenoxy group was annotated for the motif fragment *m/z* 95.0475; however, interestingly, in 34 cases this fragment was present in the MS/MS spectrum while there was no phenoxy group present in the corresponding reference molecule, nor was there any other substructure that could be assigned to this fragment. A possible explanation is that rearrangements are taking place in the mass spectrometer during the fragmentation process leading to the formation of phenoxy fragments as all these molecules do contain benzyl moieties. Here, the MAGMa-MS2LDA integration provides quick insight in assessing the consistency of structural annotations based on presence/absence of mass fragments. Furthermore, MassBank Mass2Motif 443 could be easily annotated as “aniline related” as 30 of the 32 associated molecules indeed contained an aniline or substituted aniline substructure annotated by MAGMa http://ms2lda.org/basicviz/view_parents/57561/. Finally, GNPS Mass2Motif 439 (http://ms2lda.org/basicviz/view_parents/57921/) was shown by the MAGMa annotation to originate from a specific series of oxyacetyl-amino-methyl-cyclohexane-1-carboxylic acids with a characteristic series of losses (Figure 4E). Especially with losses MAGMa annotations are very helpful during the annotation process as manual analysis of neutral losses is hampered by our inability to detect these generally smaller losses rather than larger scaffolds which are easier to recognize.

#### Chemical classification-based annotation of Mass2Motifs from standards

With increasing numbers of library MS/MS spectra available, the number of Mass2Motifs that can be extracted from those spectra will steadily increase. An alternative guidance in assigning substructures to those motifs is the use of chemical classification. We collected ClassyFire substituent terms for all the molecules in the reference MS/MS data set.^29^ These substituent terms are found using more than 5000 SMARTS patterns and are then normally used by ClassyFire to put molecules into a hierarchical chemical ontology represented by taxonomy terms. Here, we combined the substituent terms associated to molecules within a Mass2Motif to look for substituent terms that are enriched within motifs as compared to their presence across the entire data set. For example, GNPS Mass2Motif 43 was previously annotated to be related to the adenine core structure http://ms2lda.org/basicviz/view_parents/58177/. The enriched substituent terms clearly correlate with this previous annotation: terms like aminopyrimidine and 6-aminopurine are enriched as compared to the general percentage in the entire GNPS data set (Table 1). In addition, GNPS Mass2Motif 72 was enriched with amine and tertiary amine terms which is consistent with its annotation as diethylamino or dimethylaminoethyl substructure (Table 2). GNPS Mass2Motif 1 was enriched with oxosteroid related substituent terms matching its previous annotation as “sterone related” http://ms2lda.org/basicviz/view_parents/58328/. Finally, the natural product substructure of quinazolinol (4-quinazolinone) was previously assigned to GNPS Mass2Motif 60 http://ms2lda.org/basicviz/view_parents/57956/. Moreover, MAGMa annotations of MS2LDA features that were associated with this Mass2Motif also showed the quinoxaline substructure to be present in 22 out of the 25 molecules (Figure 5). The enriched ClassyFire terms confirm this annotation. This indicates that the collected terms can be used as proxy and guidance for Mass2Motif annotations in reference MS/MS data sets.

**Table 1:**
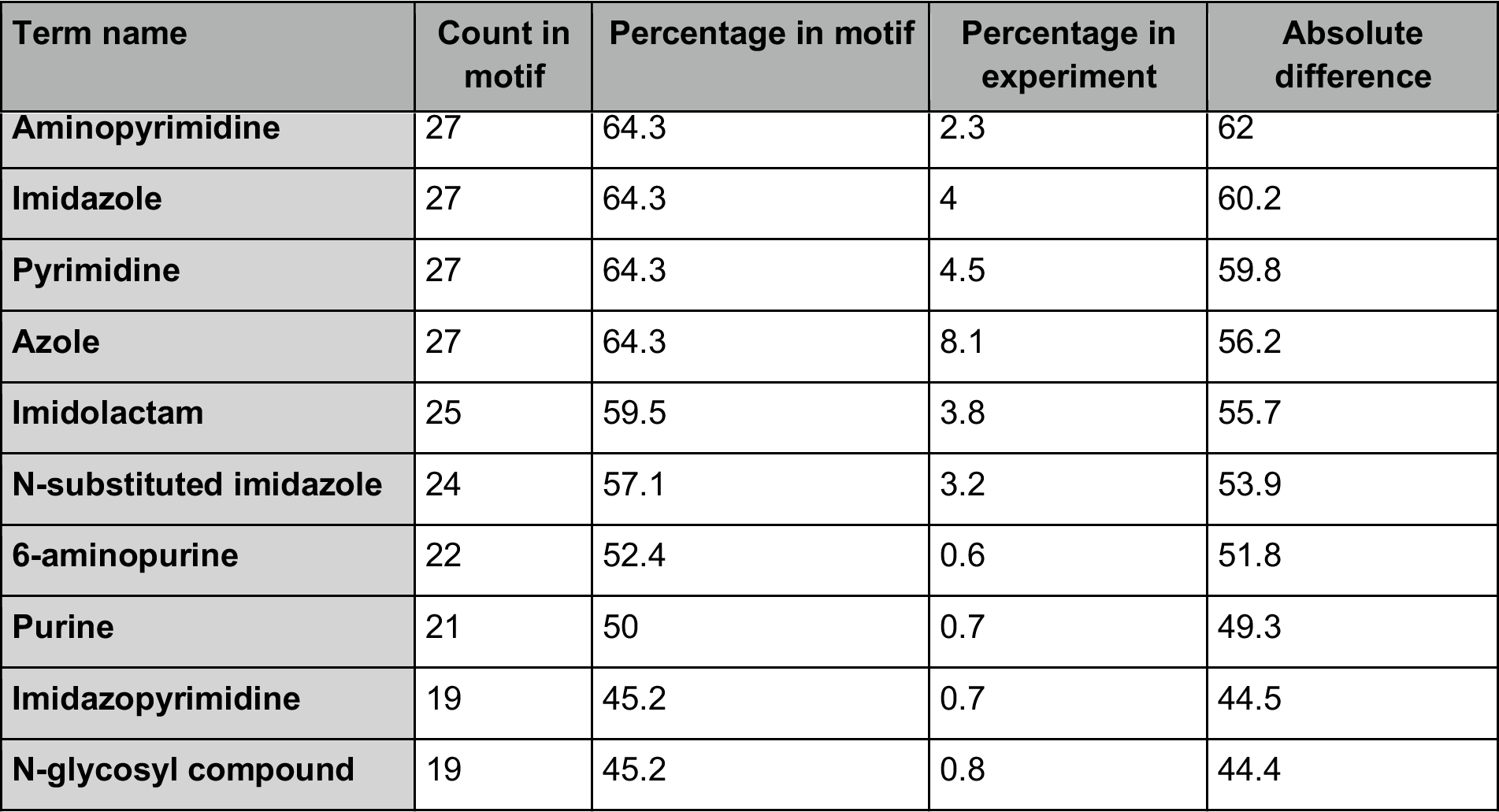
Top 10 most enriched ClassyFire substituent terms for GNPS Mass2Motif 43 previously annotated as adenine related. Term name represents the ClassyFire substituent term, Count in motif is the number of times the term appeared in a molecule associated to the Mass2Motif, Percentage in motif is the percentage of the count in motif over the total number of molecules in the motif, Percentage in experiment is the percentage of the number of term occurrences in molecules within the entire experiment over the total number of molecules, and Absolute difference is the absolute difference between the two percentages.

**Table 2:** ClassyFire substituent terms for GNPS Mass2Motif 72 annotated as diethylamino or dimethylaminoethyl substructure related. Term name represents the ClassyFire substituent term, Count in motif is the number of times the term appeared in a molecule associated to the Mass2Motif, Percentage in motif is the percentage of the count in motif over the total number of molecules in the motif, Percentage in experiment is the percentage of the number of term occurrences in molecules within the entire experiment over the total number of molecules, and Absolute difference is the absolute difference between the two percentages.

| **Term name** | **Count in motif** | **Percentage in motif** | **Percentage in experiment** | **Absolute difference** |
| --- | --- | --- | --- | --- |
| **Amine** | 49 | 58.3 | 25 | 33.3 |
| **Organoheterocyclic compound** | 6 | 7.1 | 38 | 30.9 |
| **Tertiary amine** | 38 | 45.2 | 14.6 | 30.7 |
| **Tertiary aliphatic amine** | 37 | 44 | 13.7 | 30.3 |

**Figure 5:**
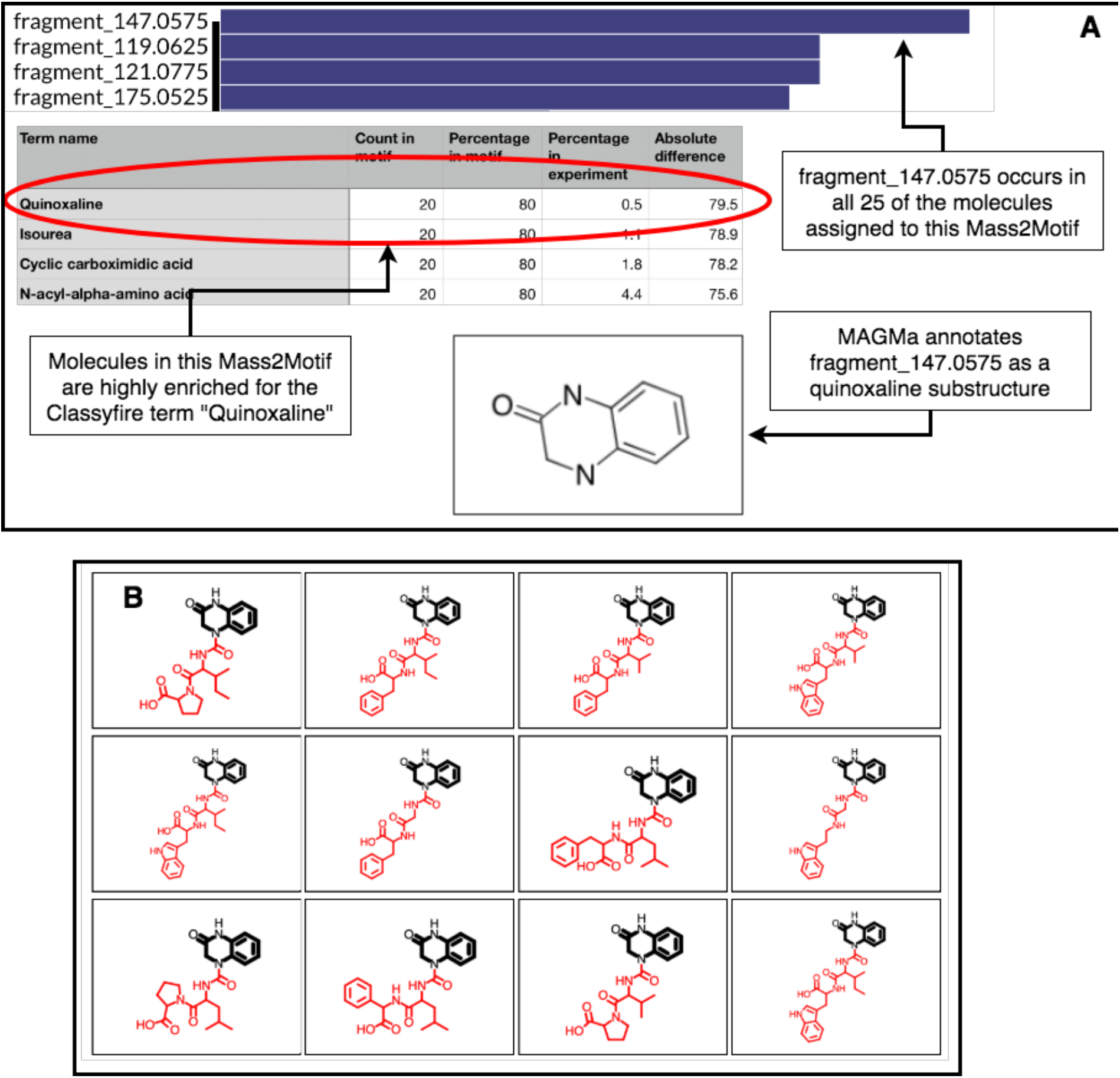
A) Top: feature frequency plot for GNPS Mass2Motif 60; Middle: Most enriched ClassyFire substituent terms in the same motif; Bottom: MAGMA assigned the quinazolinol substructure in 22 of the 25 molecules associated to this motif. B) Screenshot of the ms2lda.org web app with the by MAGMa annotated quinazolinol substructure highlighted in 12 of the 22 molecules.

With help of the new additions a number of novel annotations were made. For example, GNPS Mass2Motif 6 could be annotated with the diphenyl-containing substructure following MAGMa annotations for its mass features and its enriched ClassyFire terms http://ms2lda.org/basicviz/view_parents/58331/ (Table 3). Finally, the earlier with MAGMa annotated methoxy group related GNPS Mass2Motif 152 returned consistent ClassyFire terms being enriched in this motif such as methyl ester and carboxylic acid ester (Table 4) which is remarkable for such a small substructure. Interestingly, the with help of MAGMa annotated earlier discussed GNPS Mass2Motif 439 (Figure 4E) is not covered by helpful ClassyFire annotations, indicating the complementarity of these approaches. Overall, the enriched chemical classification terms confirmed and strengthened the manual and MAGMa annotations and as such they may support and promote the use of consistent chemical terminology during the annotation process.

**Table 3:** Top 10 most enriched ClassyFire substituent terms for GNPS Mass2Motif 6 which could in this study be annotated as diphenyl substructure related. Term name represents the ClassyFire substituent term, Count in motif is the number of times the term appeared in a molecule associated to the Mass2Motif, Percentage in motif is the percentage of the count in motif over the total number of molecules in the motif, Percentage in experiment is the percentage of the number of term occurrences in molecules within the entire experiment over the total number of molecules, and Absolute difference is the absolute difference between the two percentages.

| Term name | Count in motif | Percentage in motif | Percentage in experiment | Absolute difference |
| --- | --- | --- | --- | --- |
| Diphenylmethane | 23 | 52.3 | 2.1 | 50.2 |
| Tertiary aliphatic amine | 21 | 47.7 | 13.7 | 34 |
| Tertiary amine | 21 | 47.7 | 14.6 | 33.2 |
| Amine | 24 | 54.5 | 25 | 29.5 |
| Heteroaromatic compound | 5 | 11.4 | 36.8 | 25.4 |
| Aromatic heteropolycyclic compound | 7 | 15.9 | 40.3 | 24.4 |
| Benzenoid | 10 | 22.7 | 45 | 22.3 |
| Aromatic homomonocyclic compound | 14 | 31.8 | 9.6 | 22.2 |
| Benzylether | 8 | 18.2 | 0.6 | 17.5 |
| Dialkyl ether | 11 | 25 | 7.7 | 17.3 |

**Table 4:** Top 10 most enriched ClassyFire substituent terms for GNPS Mass2Motif 152 that was annotated with help of MAGMa as methoxy group related. Term name represents the ClassyFire substituent term, Count in motif is the number of times the term appeared in a molecule associated to the Mass2Motif, Percentage in motif is the percentage of the count in motif over the total number of molecules in the motif, Percentage in experiment is the percentage of the number of term occurrences in molecules within the entire experiment over the total number of molecules, and Absolute difference is the absolute difference between the two percentages.

| Term name | Count in motif | Percentage in motif | Percentage in experiment | Absolute difference |
| --- | --- | --- | --- | --- |
| Methyl ester | 14 | 24.1 | 2.3 | 21.8 |
| Carboxylic acid ester | 20 | 34.5 | 13.9 | 20.6 |
| Dialkyl ether | 15 | 25.9 | 7.7 | 18.2 |
| Enoate ester | 11 | 19 | 2.9 | 16.1 |
| Alpha,beta-unsaturated carboxylic ester | 11 | 19 | 2.9 | 16.1 |
| Ether | 26 | 44.8 | 30.9 | 13.9 |
| Dihydropyridinecarboxylic acid derivative | 6 | 10.3 | 0.6 | 9.8 |
| Carboxylic acid | 2 | 3.4 | 13.3 | 9.8 |
| Enamine | 5 | 8.6 | 0.6 | 8.1 |
| Monocarboxylic acid or derivatives | 16 | 27.6 | 19.7 | 7.9 |

#### Chemical classification-based annotations of Mass2Motifs from non-standards

Using more than 10,000 unique GNPS Library reference MS/MS spectra, we were able to train a neural network to infer 444 ClassyFire substituent terms from fragmentation data using 1 Da binned mass fragments as input. To evaluate how well the current model predicts enriched terms for Mass2Motifs, we performed the ClassyFire prediction analysis on a public MS2LDA experiment of 71 Rhamnaceae plant extracts (see Data Availability section). In there, more than 20 motifs were previously annotated manually.^14^ We compared the predicted terms to those manual annotations and observed the following trends for motifs predominantly based on mass fragments. For example, Rhamnaceae Mass2Motif 33 was annotated with a xylose or arabinose moiety. The ClassyFire predictions show huge enrichment of alcohol and secondary alcohol terms as well as glycosyl and O-glycosyl compounds which are all saccharide related terms http://ms2lda.org/basicviz/view_parents/109416/. The unannotated Rhamnaceae Mass2Motif 196 was enriched with overlapping terms which suggest that this is a saccharide related motif as well http://ms2lda.org/basicviz/view_parents/109504/. Rhamnaceae Mass2Motifs 3 and 86 were annotated with the 3-hydroxyflavanoid cores myricetin and quercetin, respectively http://ms2lda.org/basicviz/view_parents/109575/ and http://ms2lda.org/basicviz/view_parents/109460/. Indeed, the predicted enriched ClassyFire terms clearly point to flavonoid related terms like chromone and phenol, which is also reflective of their presence in the training data. Finally, Rhamnaceae Mass2Motif 148 was annotated as cyclopeptide alkaloids related http://ms2lda.org/basicviz/view_parents/109419/ which structures were also validated.^14^ Indeed, the predicted enriched ClassyFire terms reflect the cyclopeptide structures well with the benzene ring that is part of the cyclic structure in these alkaloid peptides (https://gnps.ucsd.edu/ProteoSAFe/gnpslibraryspectrum.jsp?SpectrumID=CCMSLIB00004679280#%7B%7D) as the most enriched term in this motif.

#### MotifDB

To demonstrate the utility of being able to match against previously annotated Mass2Motifs from MotifDB, we ran the motif matching pipeline to match newly discovered Mass2Motifs in 5021 mass spectra of a publicly available human urine sample with a set or Mass2Motifs previously manually annotated from urine samples of the same cohort run under the same experimental conditions (http://ms2lda.org/basicviz/manage_motif_matches/709/).^31^ Of the 300 Mass2Motifs discovered, 102 could be matched against 82 unique Mass2Motifs from MotifDB with cosine scores of 0.5 or greater, 41 of which had cosine scores greater than 0.9. The distribution of scores is shown in Figure 6. The ten highest scoring matches are shown in Table 5 along with the annotation, and the number of molecules that are assigned to the discovered motif (at a probability threshold of 0.1 and an overlap threshold of 0.3). In total, across the 102 motifs, a total of 3715 unique molecules include at least one of the 102 matched Mass2Motifs (out of a total of 5021 in the experiment; 74%) and 2879 (57%) unique molecules include at least one Mass2Motif matched with a score of >0.9. This percentages indicate the potential of annotating complex mixtures through substructure assignments.

**Figure 6:**
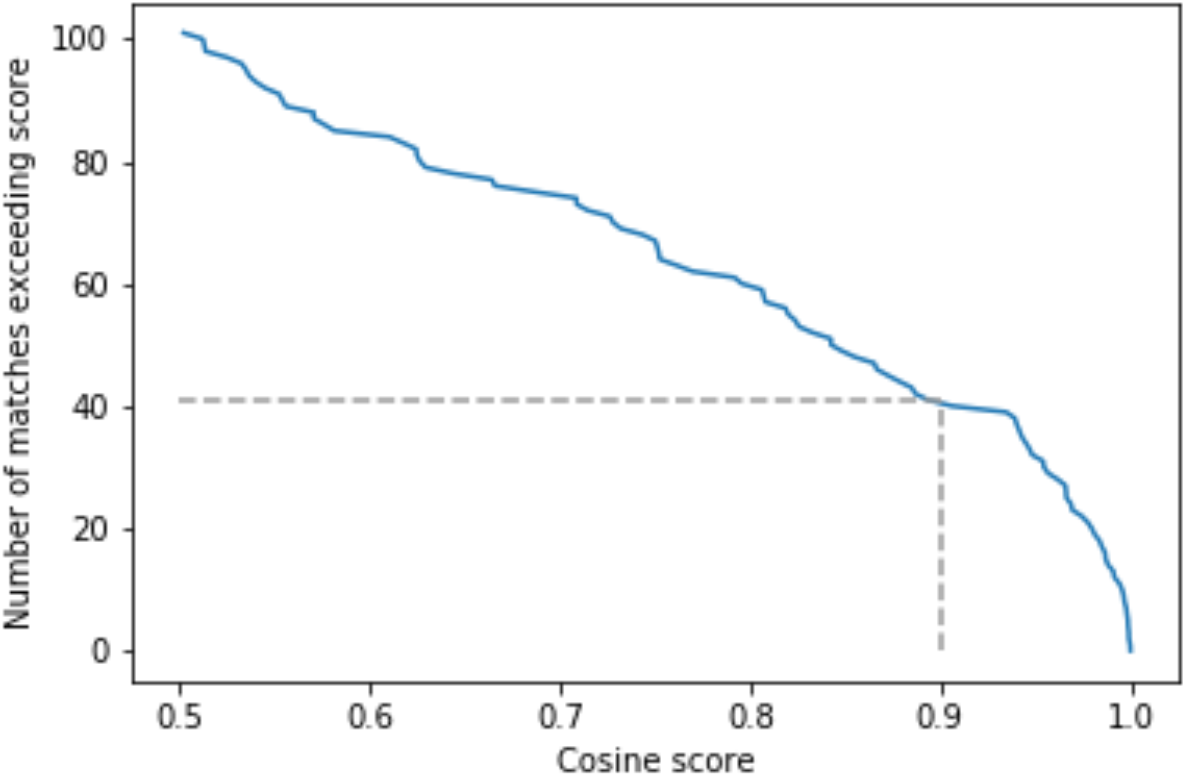
Distribution of Mass2Motif matching scores for a urine dataset matched against the urine MotifSet in MotifDB. Dashed line shows the number of Mass2Motifs (41) that could be matched against the MotifSet with a cosine score of 0.9 or more.

**Table 5:** Then highest scoring matches of Mass2Motifs discovered from urine matched against a set of urine-derived Mass2Motifs in MotifDB. Motif is the experimental Mass2Motif, MotifDB Motif is the matched motif from MotifDB, Score is the cosine similarity score between the two Mass2Motifs, MotifDB Annotation the structural annotation from MotifDB, and the Number of molecules are the number of molecules associated with the experimental Mass2Motif (out of 5021 in total).

| Motif | MotifDB Motif | Score | MotifDB Annotation | Number of molecules |
| --- | --- | --- | --- | --- |
| motif_25 | urine_mass2motif_286.m2m | 1.000 | C4H8N based Mass2Motif - indicative for proline arginine ornitine citrulline and N-containing ring structures | 217 |
| motif_181 | urine_mass2motif_233.m2m | 1.000 | Loss of 60.0248 - unclear what it points to | 86 |
| motif_49 | urine_mass2motif_194.m2m | 0.999 | Creatinine related Mass2Motif | 154 |
| motif_114 | urine_mass2motif_126.m2m | 0.999 | C7H7 and C5H5 fragments - indicative of methylbenzene substructure (aromatic) | 75 |
| motif_193 | urine_mass2motif_273.m2m | 0.999 | Fragments (C4H8NO ring fragment - and C4H7O2 and C4H5O fragments) indicative for C4H10NO2 amino acid substructure | 40 |
| motif_206 | urine_mass2motif_230.m2m | 0.999 | Water loss - indicative of a free hydroxyl group | 147 |
| motif_238 | urine_mass2motif_18.m2m | 0.999 | C2H3N loss - could be specific for a type of ring? | 130 |
| motif_151 | urine_mass2motif_293.m2m | 0.998 | Carnitine related Mass2Motif ,acylcarnitines are prevalent | 419 |
| motif_68 | urine_mass2motif_228.m2m | 0.997 | Lysine related Mass2Motif | 191 |
| motif_72 | urine_mass2motif_27.m2m | 0.996 | C5H10NO and C3H6N fragments - most likely N-methyl-morpholine substructure | 138 |

To further evaluate the power of motif matching against MotifDB we matched the urine motif set from MotifDB with Mass2Motifs discovered in fragmentation spectra of 6 urines from another cohort analysed under the same experimental conditions (http://ms2lda.org/basicviz/manage_motif_matches/601/).^21^ In this case, of the 200 Mass2Motifs discovered in the experiment, 55 could be matched at a threshold of at least 0.5 (covering 573 of the 1163 molecules; 49%) and 20 at a threshold of 0.9 (404 molecules; 35%). Although, as expected since the data is from a different cohort, the number of matches is lower than that for the first example, the ability to immediately match approximately a quarter of the discovered motifs (allowing some level of annotation for half of the molecules) highlights the generalizability of Mass2Motifs across sample sets, and the power of matching against previously discovered ones. This approach also aids in the discovery and prioritization of novel Mass2Motifs that may well represent xenobiotic-related chemistry (i.e., drug, food, etc.) not previously encountered.

## Conclusions and Future Outlook

In this paper, we have described multiple extensions to the MS2LDA platform (all implemented at the ms2lda.org web app) that enhance the ability of analysts to characterize the makeup of complex mixtures of metabolites. The extensions all make it easier to characterize the Mass2Motifs onto which MS2LDA allows experimental data to be decomposed. These Mass2Motifs often represent chemical substructures and annotating them allows some degree of annotation to all MS2 spectra that include them as often a relative small number of annotated Mass2Motifs provide information about a significant proportion of the molecules in an experiment.^8^

The extensions move the platform forward in two general directions. The first, MotifDB, provides a platform that allows for the storage of annotated Mass2Motifs that can then be accessed via an API (details at http://ms2lda.org/motifdb) or used within ms2lda.org by allowing users to match Mass2Motifs discovered within their experiments to those stored in MotifDB. In our experiments with human urine data, we found that roughly 25% of the Mass2Motifs in a urine dataset from a different cohort than the dataset from which the annotated motifs were generated could be matched against Mass2Motifs from MotifDB. These 25% of Mass2Motifs were associated to about 50% of the molecules.

The second direction is the collation of known and predicted molecular properties for individual molecules across Mass2Motifs. Here, we have presented three advances. Firstly, the use of MAGMa on databases of standards that had been analysed with MS2LDA to annotate their fragment spectra with substructures. We show how MAGMa-Mass2Motif annotations provide quick insight in ambiguity of annotations in case of isomeric substructures. These substructure annotations can then be propagated to the features in the Mass2Motifs, providing relevant insight into the substructures they could represent.

The second advance propagates the Classyfire substituent terms for the same standards datasets to the Mass2Motif level. Finally, for “unknown” molecules from experimental data of mixtures, we have introduced a machine learning approach based on a neural network that can predict a subset of classyfire substituent terms from the spectral data. The current implementation has some limitations: i) the predictive power is dependent on the chemical diversity present in available training spectra, ii) the current training set consists of series of structurally correlated molecules, and iii) very small substructures will be difficult to predict due to their usually widespread presence which makes it harder to recognize specific chemical terms in the diverse molecules. Nevertheless, we show that for fragment-based Mass2Motifs from complex mixtures the predicted terms can guide Mass2Motif annotations. Again, these can be propagated to the Mass2Motif level, providing insight into their structural makeup. We foresee that by annotating more and more Mass2Motifs, the metabolite annotation of yet unknown molecules in complex mixtures - the main bottleneck in untargeted metabolomics data analysis - will become easier. Here, we show that the neutral networking approach has the potential for further exploration and optimization. This is an avenue for future work - the model can be further augmented by inclusion of neutral loss features as well as mass shift features which is expected to improve chemical predictions for loss-based motifs such as loss of hexose or deoxyhexose and amino acid related motifs, respectively.

In the future, we anticipate these tools becoming even more useful. As more Mass2Motifs are extracted and annotated from the growing datasets of standards, MotifDB will grow and the coverage across experiments will increase. We also foresee that instead of unsupervised discovery of fragmentation patterns alone, users can include annotated motif sets of their choice to their LDA experiment thereby both finding known substructure patterns and discovering yet unknown ones. Such a workflow can then replace the current “Decomposition” workflow that only uses defined annotated motif sets and has the benefit of combining supervised and unsupervised motif discovery in one analysis. Furthermore, users would then also be able to decompose single spectra over these motif sets through an API.

The MAGMa and ClassyFire based annotations can significantly enhance the process of annotation of the rapidly growing (number of) datasets and Mass2Motifs. The expected growth in available fully annotated reference spectra will also increase the training sets available for our ClassyFire predictor, increasing performance and increasing the set of terms that we can confidently predict. Furthermore, the implementation of chemical ontology from ClassyFire assists in more consistent annotations of motifs by using chemical terminology from an ontology.

We expect that substructure-based annotation strategies will prove to be essential to decipher complex mixtures and enable meaningful biochemical interpretation. Our work represents key steps of this workflow by recognizing mass spectral patterns, semi-automated structurally annotating and storing them. An increasing amount of structurally annotated Mass2Motifs will allow metabolomics researchers to gain some structural information on the majority of fragmented molecules. The further closing of the structural annotation gap in metabolomics will make untargeted metabolomics a very powerful tool to study complex mixtures.

## Author contributions

SR, LR, and JJJvdH conceptualized the study. LR and JW designed and implemented MAGMa-MS2LDA integration. ME extracted well-annotated publicly available spectral data from GNPS. SR and JJJvdH designed ClassyFire predictions. CWO and SR built the neural network model for ClassyFire predictions and SR integrated it within MS2LDA. CWO, LR, SR, and JJJvdH analyzed data. All authors contributed to the writing of the manuscript and agreed on the content.

## Acknowledgements

The authors would like to thank all GNPS contributors who took the efforts to extensively annotate their public spectra including SMILES which made them reusable in this study. The authors would also like to thank the ClassyFire initiative for sharing the chemical ontology with the scientific community.

